## Supplementary Material for "Bacterial microbiota composition of *Ixodes ricinus* ticks: the role of environmental variation, tick characteristics and microbial interactions"

### Variables selection and justification

Tick activity is influenced by abiotic factors, such as temperature and humidity (Gray 1991, Gern et al. 2008) Using the central coordinate for each sampling site, we therefore calculated the number of days in 2014 in which the temperature was > 7 C°, as this has been suggested to be the threshold for tick activity (Lindgren et al. 2000, Perret et al. 2000). High temperatures can break tick developmental diapause and, in general, higher temperatures speed up their life cycle (Gardiner et al. 1981). Furthermore, dry conditions during summer can cause mortality in diapausing ticks (Randolph 2001, Gray 2008). For each site, we therefore calculated monthly precipitation and average temperature. Precipitation may be an inaccurate proxy for the microclimatic conditions experienced by ticks (Alonso-Carné et al. 2016), yet it has been shown to correlate with tick activity patterns (Barandika et al. 2006, Cat et al. 2017). Finally, vegetation type can affect tick abundance (Hoch et al. 2010, Tack et al. 2012, Estrada-Peña et al. 2016) and tick-borne pathogen occurrence (James et al. 2014, Raghavan et al. 2016). We therefore determined the proportion of forest cover in a 500 m radius for each sampling site. To quantify topography, aspect values were converted so that 0 corresponds to south, 90 to west and east and 180 to north.

At each sampling site, we obtained information on *I. ricinus* abundance from Lemoine et al. (2018), as it has been suggested that higher tick densities may be associated with higher tick-borne pathogen prevalence (Jouda et al. 2004, Walk et al. 2009). We also obtained information on the abundance of a key tick host at the sampling sites, the bank vole (*Myodes glareolus*), as well as the ratio of bank vole to other rodents from Cornetti et al. (2018). Bank voles are particularly competent hosts for pathogens such as *Borrelia afzelii*, whereas other rodents have been suggested to be less competent (Hanincova et al. 2003). The relative abundance of bank voles and other rodent may thus influence the prevalence of pathogens because of 'dilution effects' (Kurtenbach et al. 1998, LoGiudice et al. 2003, Keesing et al. 2006, Begon 2008).

The genetic make-up of the host may affect pathogen and endosymbiont colonisation and replication success (Archie and Ezenwa 2011). In order to quantify individual and population-level genetic diversity, we genotyped ticks at 11 microsatellite markers in two multiplexed amplifications. The first multiplex panel consisted of the primer pairs IRN7, IR39, IRic13, IRic11, IRic08, IRic09 and IRN31, whereas the second multiplex panel consisted of the primers IRN37, IRN12, IRic18 and IRic05 (Delaye et al. 1998, Røed et al. 2006, Kempf et al. 2011). Amplification took place in a total volume of 6 µl, including 3 µl of 2x Qiagen PCR Master Mix (Qiagen; Hilden, Germany), 1.5 µl of tick DNA, 1 µl of 10 x primer mix and 0.5 µl of $H_20$ per sample. The PCR protocol consisted of an initial denaturation step for 15 minutes at 95 C°, followed by 35 cycles of 30 seconds at 95 C°, 90 seconds at 58 C° and 60 seconds at 72 C° and a final elongation step at 60 C° for 30 minutes for both multiplex panels. The amplified products were diluted 1:20 and 1 µl of the diluted product was mixed with 18 µl of HiDi-LIZ (Applied Biosystems, Foster City, CA, USA). Sequencing was performed on an ABI Prism 3730 capillary sequencer (Applied Biosystems, Foster City, CA, USA). Fragment length was determined using Genemapper 3.7 (Applied Biosystems, Foster City, CA, USA).

**Table S1:** Model input variables, their assignment to different components of the model and the level at which each variable has been measured.

| Component | | Variable | Level |
|---|---|---|---|
| Occurrence | | Presence-absence | Individual / OTU |
| Traits | | Endosymbiont | OTU |
| | | Human or wildlife pathogen | OTU |
| Spatial context | | Latitude | Site |
| | | Longitude | Site |
| Environment | Full variable set | Intercept | Individual |
| | | Tick sex | Individual |
| | | Tick life stage | Individual |
| | | Tick heterozygosity | Individual |
| | | Tick abundance | Site |
| | | Elevation | Site |
| | Variable selection set | Expected tick population heterozygosity | Site |
| | | Number of days > 7 C° during the year | Site |
| | | Monthly precipitation | Site / Month |
| | | Mean monthly temperature | Site / Month |
| | | Forest coverage (500 m radius | Site |
| | | Slope | Site |
| | | Aspect | Site |
| | | Vole abundance | Site |
| | | Proportion of voles to other rodents | Site |
| Latent variables (random effects) | | Tick ID | |
| | | Location | |
| | | Site | |
| | | Month | |

Tick microbiota sequencing

16S sequencing libraries were prepared following the Earth Microbiome 16S Illumina Amplicon protocol, using the primers

515FB: CTTTCCCTACACGACGCTCTTCCGATCTNNNNNNNNGTGYCAGCMGCCGCGGTAA
806RB: GGAGTTCAGACGTGTGCTCTTCCGATCTNNNNNNNNGGACTACNVGGGTWTCTAAT

which also contained the Illumina Miseq primer binding site (Caporaso et al. 2012, Apprill et al. 2015, Walters et al. 2015). Samples and negative controls were randomized across two plates. Each reaction contained 1 µl of template DNA with 0.5 µM forward and reverse primers, 0.3 mM dNTP, 0.4 µl KAPA HiFi HotStart polymerase (KAPA Biosystems, Basel, Switzerland) in a final volume of 20 µl. The PCR protocol started with an initial denaturation at 95 C° for 3 minutes, followed by 35 cycles of 20 seconds at 98 C°, 60 seconds at 50

C° and 30 seconds at 72 C° with final elongation step of 5 minutes at 72 C°. We ran PCR products on 2% agarose gels, cut out the bands corresponding to the amplicon size (350 bp) and eluted them in 10 µl milli-Q water. We then purified the extracted amplicons using MinElute Gel Extraction kit (Qiagen, Hilden, Germany) following the manufacturer's protocol. In second PCR step Illumina Miseq adaptors and dual indexing Miseq barcodes were incorporated into the previously amplified V3-V4 regions. The second PCR protocol started with an initial denaturation at 95 C° for 3 minutes, followed by 15 cycles of 20 seconds at 98 C°, 60 seconds at 54 C° and 30 seconds at 72 C° with final elongation step of 5 minutes at 72 C°. Products were visualized and purified as described above. We then measured the DNA concentration of the purified product with Invitrogen Qubit 4 Fluorometer ssDNA assay kit (Thermo Fisher Scientific, Walthan, MA, USA) and created library pools by mixing equimolar amounts of purified products to reach a 4 nM library concentration. The libraries were further processed using standard normalization protocols with PhiX controls and sequenced on a Illumina MiSeq at the Functional Genomic Center Zurich using reagent kit 600cycle v3 (Illumina, San Diego, CA, USA) with a target length of 250 bp per amplicon following the manufacturer's protocol.

*mothur* analysis and oligotyping

Using *mothur*, we purged unsuccessful contigs and preserved only contigs between 250 and 310 bp. The alignment was made against aligned SILVA bacterial references (release 128; https://www.arb-silva.de/documentation/release-128/). We used 97% similarity to determine OTUs and classified them with the *wang* method (Wang et al. 2007) using SILVA taxonomy.

For oligotyping analysis, we did initial entropy analysis, and if unexplained entropy was found, we oligotyped with the 2 or 3 highest entropy positions, depending on the results of the entropy analysis. After this we repeated oligotyping until the oligotypes converged, that is, there was no formation of new oligotypes observed when existing oligotypes were further decomposed.

We identified endosymbionts (i.e., any bacteria which live within tick cells and have positive, neutral or negative effects on their hosts) and human or wildlife pathogens (i.e., any bacteria that are (or have close relatives that are) pathogenic to vertebrates) in the OTU data. OTUs were classified as endosymbionts, pathogens, both, or neither, based on a literature search for each taxonomic label (Table 2 in main text) in Web of Science (Clarivate Analytics, Philadelphia, PA, USA) using the taxonomic label and 'pathogen' or 'endosymbiont' as search terms. The following references were used for OTU assignment: *Midichloria* endosymbiont (Sassera et al. 2006), *Spiroplasma* endosymbiont (Tully et al. 1981), *Rickettsiella* endosymbiont (Kurtti et al. 2002), *Lariskella* endosymbiont (Matsuura et al. 2012), *Rickettsia helvetica* pathogen and endosymbiont (Beati et al. 1993), *R. monacensis* pathogen and endosymbiont (Sekeyova et al. 2000), *Rickettsia sp.* pathogen and endosymbiont (Perlman et al. 2006), *Anaplasma* pathogen and endosymbiont (Stuen 2007), *Candidatus* Neoehrlichia pathogen (Kawahara et al. 2004), *Borrelia afzelii* pathogen (Marin Canica et al. 1993), *B. miyamotoi* pathogen (Fukunaga et al. 1995), *B. garinii* pathogen (Baranton et al. 1992) and *B.*

*valaisiana* pathogen (Wang et al. 1997). The pathogens identified in our study have previously been described in *I. ricinus* in Switzerland and Central Europe (Lommano et al. 2012, Strnad et al. 2014, Oechslin et al. 2017).

Hierarchical modelling of Species Communities

We used a Hierarchical Modelling of Species Communities (HMSC) approach to model tick-associated bacterial microbiota. The input data for this model is a matrix of species occurrence and a matrix of environmental variables. Additional data included in the model consist of species traits to model species niches and spatiotemporal context of sampled species (Figure S1). This modelling approach provides estimates for species niches, which are the responses of OTUs to tick-related and environmental variables. Estimates of variation in (co-)occurrence can be used to infer species-to-species associations, when taking shared environmental conditions into account.

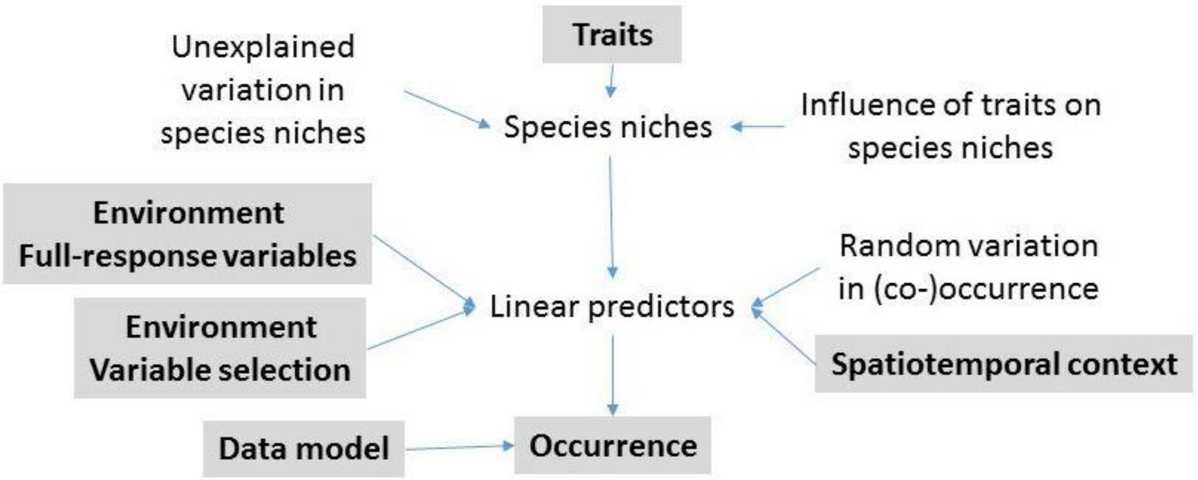


**Figure S1**: Hierarchical Modelling of Species Communities model structure. Components in gray are the input variables. Modified from Ovaskainen et al. 2017b.

Our samples are hierarchically structured as there are three sampling locations each containing three sites at different elevations. Furthermore, the samples have been collected during three months (June, July, August). Consequently, the environmental and spatial context variables have been measured at different levels: while tick-related variables are generally related to the individual samples, i.e., each sample has its own value, the population and topographic variables, such as population genetic structure and slope, are measured at a level of site, and climate variables such as temperature and precipitation are measured at the level of both site and month (Table S1).

Environmental variables were split in two different variable sets: full-effect variables were used in the model in every iteration, whereas for the variable selection variables only a subset was included in each iteration. Variable selection assigns an indicator value for each variable which is either 0 or 1. If the indicator value is 0, the variable is not included in the iteration, whereas if the value is 1, it is included. Indicator values were determined from full conditional distributions by computing the likelihood of the data for both values 0 and 1. We used the default priors of the framework for full-effect variables and a uniform prior of 0.1 for all variables in the variable selection set (Ovaskainen et al. 2017a).

A Bernoulli distribution with a probit link function was used to model the response community data matrices with Bayesian inference. We ran 100 000 Markov chain Monte Carlo (MCMC) iterations, out of which the first 70 000 were discarded, with the remaining thinned to 620 posterior samples. The model was fitted using MATLAB R2017a (MathWorks Inc., Natick, MA, USA). We sampled the posterior distribution using the Gibbs sampler (Ovaskainen et al. 2017b) and we checked for model convergence by visually assessing trace plots.

We used block cross-validation for mode fit assessment (Roberts et al. 2017). We selected randomly 10 samples, including at least one sample from each site, using the rest of the samples (N = 72) as the training set. We assessed the model fit by predicting the validation data with the training model and comparing it to the true occurrences by calculating the Tjur $R^2$ coefficients of discrimination (Tjur 2009) for each OTU. Furthermore, as a site-level coefficient of discrimination we calculated Spearman’s rank correlations between the predicted and true occurrences at the levels of sampling units and sites. A good model fit was observed with a mean coefficient of discrimination for five iterations of cross-validation of 0.18 (range: 0.14-0.20) for tick ID level and > 0.90 (range: 0.91-0.99) for the site level.

Variance partitioning assesses the explanatory power of different (groups of) explanatory variables in relation to the same response variable and thus can reveal which environmental variables are more or less influential (Borcard et al. 1992). We partitioned the variation explained by the fixed effects (i.e., the variables included in the full-effect and the variable selection sets) and by random effects (i.e., the study design). Within the latter, we further separated the variation explained by month, sampling site, location, and tick ID.

Using a variance partitioning approach, we found that tick-level fixed effects (i.e., sex, life stage, genetic diversity) played a particularly important role in explaining variation in the occurrence of pathogens and endosymbionts across ticks, explaining on average 17.5% of the total variation (Figure 2 in main text). This proportion increased to 23.9% for *Candidatus* Neoehrlichia, the highest proportion explained by tick-related fixed effects for any OTU (Figure 2 in main text). Furthermore, the environmental variables elevation and temperature played a particularly important role in explaining variation in the occurrence of pathogens and endosymbionts across ticks, explaining on average 14.3% of the total variation (Figure 2 in main text).

Tick sex / life stage influenced the occurrence of pathogens, with adult females being more likely to harbour pathogens (Figure S2) than males. These variables explained 11.4 % of the total variation explained by the model.

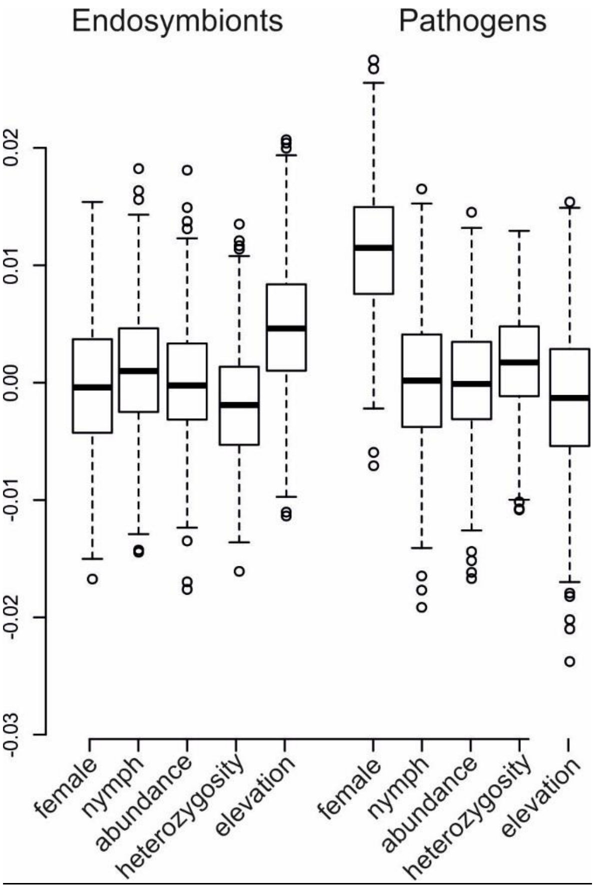


**Figure S2**: The effects of full-variable set variables on the occurrence of endosymbiont or pathogen OTUs. The only strongly supported association was a higher probability of pathogen occurrence in females. Positive values indicated a higher probability of occurrence and negative values a lower probability. Boxplots show interquartile range with vertical lines showing highest and lowest data points still within 1.5 interquartile range from upper or lower quartile.

We estimated effect sizes of the different environmental variables by using the full model to predict interpolated values for one variable at a time, while keeping the other variables constant. Specifically, for each OTU and variable which had strong statistical support, we estimated effect sizes by predicting 100 values evenly spaced over the whole range of actual values in the original model while setting other environmental variables to mean, latent variables to a single sampling unit, location and month outside of the actual bounds and site and geographical coordinates to the most central of our sites.

The effect sizes varied substantially across OTUs and explanatory variables (Figure S3a-i). For example a threefold increase in tick abundance was associated with a threefold increase in *Neoehrlichia* prevalence from 8% to 26% (Figure S3e). Other large effect sizes included a tick sex differences in *Lariskella* (females with mean 49%±4% based on the 90% central credible interval, males 62%±5%), *Rickettsiella* (females 54%±4%, males 79%±4%) and *Spiroplasma* (females 29%±3%, males 40% ±4%) prevalence, differences in *B. garinii* (from 12%± 32% at 650 masl to 2% ±15% at 1500 masl), *R. helvetica* (from 12%±33% at 650 masl to 37%±49% at 1500 masl) and *R. monascensis* (from 5% ± 22% at 650 masl to 23% ± 40% at 1500 masl) prevalence along elevational clines, the change in *Lariskella* prevalence (from 66% ±47% at 0.2 to 43% ±49% at 1.0) with tick individual-level heterozygosity and the association between aspect and *Rickettsiella* occurrence (from 73% ±45% at southfacing sites to 55%±50% at northfacing sites) and aspect and Borrelia afzelii occurrence (from 5% ±21% at southfacing sites to 27%±44% at northfacing sites).

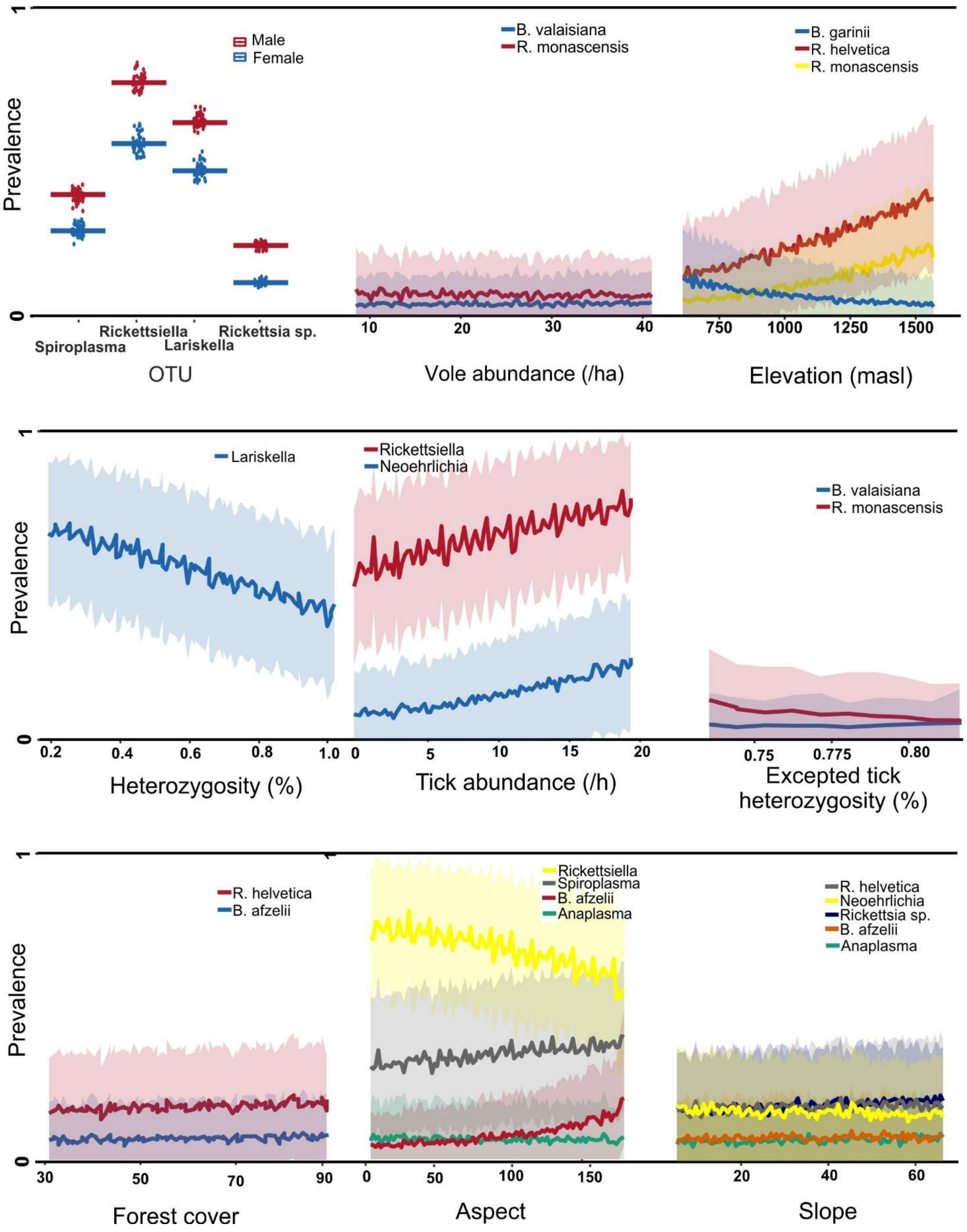


**Figure S3**: Occurrence of endosymbionts and pathogens in ticks along environmental gradients. The predictions of all explanatory variables with strong statistical support are shown. We used the full model to predict interpolated values for one variable at a time for 100 evenly spaced values within the observed range of actual values, while keeping the other variables constant and set to their mean value for the illustration of focal variable effect. The error represents the 90% central credible interval.

**Table S2:** Co-occurrence of endosymbionts and pathogens within ticks after accounting for shared environmental preferences. Positive values represent the strength of association when OTUs are more probable to co-occur and negative values when OTUs are less probable to co-occur than expected by chance. Only associations with strong statistical support (based on the 90% central credible interval) are presented.

| | | *Spiroplasma* | *Rickettsiella* | *Lariskella* | *Rickettsia helvetica* | *R. monacensis* | *Rickettsia sp.* | *Anaplasma* | *Ca.* Neoehrlichia | *Borrelia afzelii* | *B. miyamotoi* | *B. garinii* | *B. valaisiana* |
|---|---|---|---|---|---|---|---|---|---|---|---|---|---|
| Otu0003 | *Spiroplasma* | | | | | | | | | | | | |
| Otu0005 | *Rickettsiella* | | | | | | | | | | | | |
| Otu0022 | *Lariskella* | -0.93 | | | | | | | | | | | |
| Otu0031 | *Rickettsia helvetica* | | 0.70 | | | | | | | | | | |
| | *R. monacensis* | | | | | | | | | | | | |
| Otu0067 | *Rickettsia sp.* | -0.90 | | 0.90 | | | | | | | | | |
| Otu0076 | *Anaplasma* | | | | | | | | | | | | |
| Otu0086 | *Ca.* Neoehrlichia | -0.34 | 0.68 | | | | 0.85 | | | | | | |
| Otu0088 | *Borrelia afzelii* | | 0.61 | | 0.63 | | | | | | | | |
| | *B. miyamotoi* | -0.38 | | 0.43 | | | | | | | | | |
| | *B. garinii* | | 0.61 | | 0.69 | | 0.41 | | 0.64 | 0.61 | | | |
| | *B. valaisiana* | | 0.82 | | 0.65 | | | | | 0.61 | | 0.65 | |

### *Supplementary references*


Alonso-Carné, J. et al. 2016. Modelling the phenological relationships of questing immature Ixodes ricinus (Ixodidae) using temperature and NDVI Data. - Zoonoses Public Health 63: 40–52.

Apprill, A. et al. 2015. Minor revision to V4 region SSU rRNA 806R gene primer greatly increases detection of SAR11 bacterioplankton. - Aquat. Microb. Ecol. 75: 129–137.

Archie, E. a and Ezenwa, V. O. 2011. Population genetic structure and history of a generalist parasite infecting multiple sympatric host species. - Int. J. Parasitol. 41: 89–98.

Barandika, J. F. et al. 2006. Risk factors associated with ixodid tick species distributions in the Basque region in Spain. - Med. Vet. Entomol. 20: 177–188.

Baranton, G. et al. 1992. Delineation of Borrelia burgdorferi Sensu Stricto, Borrelia garinii sp . nov ., and Group VS461 associated with Lyme Borreliosis. - Int. J. Syst. Bacteriol. 42: 378–383.

Beati, L. et al. 1993. Species of the Spotted Fever Group of Rickettsiae. - Int. J. Syst. Bacteriol. 43: 521–526.

Begon, M. 2008. Effects of host diversity on disease dynamics. - In: Ostfeld, R. S. et al. (eds), Infectious Disease Ecology: Effects of Ecosystems on Disease and of Disease on Ecosystems. Princeton University Press, pp. 12–29.

Borcard, D. et al. 1992. Partialling out the spatial component of ecological variation. - Ecology 73: 1045–1055.

Caporaso, J. G. et al. 2012. Ultra-high-throughput microbial community analysis on the Illumina HiSeq and MiSeq platforms. - ISME J. 6: 1621–1624.

Cat, J. et al. 2017. Influence of the spatial heterogeneity in tick abundance in the modeling of the seasonal activity of Ixodes ricinus nymphs in Western Europe. - Exp. Appl. Acarol. 71: 115–130.

Cornetti, L. et al. 2018. Small-scale spatial variation in infection risk shapes the evolution of a *Borrelia* resistance gene in wild rodents. - Mol. Ecol.: 3515–3524.

Delaye, C. et al. 1998. Isolation and characterization of microsatellite markers in the Ixodes ricinus complex (Acari: Ixodidae). - Mol. Ecol. 7: 357–363.

Estrada-Peña, A. et al. 2016. Perspectives on modelling the distribution of ticks for large areas: so far so good? - Parasit. Vectors 9: 179.

Fukunaga, M. et al. 1995. Genetic and phenotypic analysis of Borrelia miyamotoi sp. nov., isolated from the ixodid tick Ixodes persulcatus, the vector for Lyme Disease in Japan. - Int. J. Syst. Bacteriol. 45: 804–810.

Gardiner, W. P. et al. 1981. Models based on weather for the development phases of the sheep tick, Ixodes ricinus L. - Vet. Parasitol. 9: 75–86.

Gern, L. et al. 2008. Influence of some climatic factors on Ixodes ricinus ticks studied along altitudinal gradients in two geographic regions in Switzerland. - Int. J. Med. Microbiol. 298: 55–59.

Gray, J. S. 1991. The development and seasonal activity of the tick Ixodes ricinus: a vector of Lyme borreliosis. - Rev. Med. Vet. Entomol. 79: 323–333.

Gray, J. S. 2008. Ixodes ricinus seasonal activity: Implications of global warming indicated by revisiting tick and weather data. - Int. J. Med. Microbiol. 298: 19–24.

Hanincova, K. et al. 2003. Association of Borrelia afzelii with rodents in Europe. - Parasitology 126: 11–20.

Hoch, T. et al. 2010. Influence of host migration between woodland and pasture on the population dynamics of the tick Ixodes ricinus: A modelling approach. - Ecol. Modell. 221: 1798–1806.

James, M. C. et al. 2014. The heterogeneity,distribution, and environmental associations of Borrelia burgdorferi Sensu Lato, the agent of Lyme borreliosis, in scotland. - Front. Public Heal. 2: 1–10.

Jouda, F. et al. 2004. Ixodes ricinus density, and distribution and prevalence of Borrelia burgdorferi sensu lato infection along an altitudinal gradient. - J. Med. Entomol. 41: 162–169.

Kawahara, M. et al. 2004. Ultrastructure and phylogenetic analysis of "Candidatus Neoehrlichia mikurensis" in the family Anaplasmataceae, isolated from wild rats and found in Ixodes ovatus ticks. - Int. J. Syst. Evol. Microbiol. 54: 1837–1843.

Keesing, F. et al. 2006. Effects of species diversity on disease risk. - Ecol. Lett. 9: 485–98.

Kempf, F. et al. 2011. Host races in Ixodes ricinus, the European vector of Lyme borreliosis. - Infect. Genet. Evol. 11: 2043–2048.

Kurtenbach, K. et al. 1998. Differential transmission of the genospecies of Borrelia burgdorferi sensu lato by game birds and small rodents in England. 64: 1169–1174.

Kurtti, T. J. et al. 2002. Rickettsiella -like Bacteria in Ixodes woodi ( Acari : Ixodidae ).: 534–540.

Lindgren, E. et al. 2000. Impact of climatic change on the northern latitude limit and population density of the disease-transmitting European tick Ixodes ricinus. - Environ. Health Perspect. 108: 119–123.

LoGiudice, K. et al. 2003. The ecology of infectious disease: Effects of host diversity and community composition on Lyme disease risk. - Proc. Natl. Acad. Sci. 100: 567–571.

Lommano, E. et al. 2012. Infections and coinfections of questing Ixodes ricinus ticks by emerging zoonotic pathogens in Western Switzerland. - Appl. Environ. Microbiol. 78: 4606–4612.

Marin Canica, M. et al. 1993. Monoclonal antibodies for identification of Borrelia afzelii sp. nov. associated with late cutaneous manifestations of Lyme Borreliosis. - Scand. J. Infect. Dis. 25: 441–448.

Matsuura, Y. et al. 2012. Novel clade of alphaproteobacterial endosymbionts associated with stinkbugs and other arthropods. - Appl. Environ. Microbiol. 78: 4149–4156.

Oechslin, C. P. et al. 2017. Prevalence of tick-borne pathogens in questing Ixodes ricinus ticks in urban and suburban areas of Switzerland. - Parasites and Vectors 10: 1–18.

Ovaskainen, O. et al. 2017a. How are species interactions structured in species-rich communities ? A new method for analysing time-series data. - Proc. R. Soc. B Biol. Sci. 284: 20170768.

Ovaskainen, O. et al. 2017b. How to make more out of community data? A conceptual framework and its implementation as models and software. - Ecol. Lett. 20: 561–576.

Perlman, S. J. et al. 2006. The emerging diversity of Rickettsia. - Proc. R. Soc. B Biol. Sci. 273: 2097–2106.

Perret, J.-L. et al. 2000. Influence of saturation deficit and temperature on Ixodes ricinus tick questing activity in a Lyme borreliosis-endemic area (Switzerland). - Parasitol. Res. 86: 554–557.

Raghavan, R. K. et al. 2016. Heterogeneous associations of ecological attributes with tick-borne rickettsial pathogens in a periurban landscape. - Vector-Borne Zoonotic Dis. 16: 569–576.

Randolph, S. E. 2001. The shifting landscape of tick-borne zoonoses: tick-borne encephalitis and Lyme borreliosis in Europe. - Philos. Trans. R. Soc. London B Biol. Sci. 356: 1045–56.

Roberts, D. R. et al. 2017. Cross-validation strategies for data with temporal, spatial, hierarchical, or phylogenetic structure. - Ecography (Cop.). 40: 913–929.

Røed, K. H. et al. 2006. Identification and characterization of 17 microsatellite primers for the tick, Ixodes ricinus, using enriched genomic libraries. - Mol. Ecol. Notes 6: 1165–1167.

Sassera, D. et al. 2006. Candidatus Midichloria mitochondrii, an endosymbiont of the Ixodes ricinus with a unique intramitochondrial lifestyle. - Int. J. Syst. Evol. Microbiol. 56: 2535–2540.

Sekeyova, Z. et al. 2000. Characterization of a new spotted fever group rickettsia detected in Ixodes ricinus (Acari: Ixodidae) collected in Slovakia. - J Med Entomol. 37: 707–713.

Strnad, M. et al. 2014. Europe-wide meta-analysis of Borrelia burgdorferi sensu lato prevalence in questing Ixodes ricinus ticks. - Appl. Environ. Microbiol. in press.

Stuen, S. 2007. Anaplasma Phagocytophilum - The most widespread tick-borne infection in animals in Europe. - Vet. Res. Commun. 31: 79–84.

Tack, W. et al. 2012. Local habitat and landscape affect Ixodes ricinus tick abundances in forests on poor, sandy soils. - For. Ecol. Manage. 265: 30–36.

Tjur, T. 2009. Coefficients of determination in logistic regression models — a new Proposal: the coefficient of fiscrimination. - Am. Stat. 63: 366–372.

Tully, J. G. et al. 1981. Helical mycoplasmas (Spiroplasmas) from Ixodes ticks. - Science (80-. ). 212: 1043–1046.

Walk, S. T. et al. 2009. Correlation between tick density and pathogen endemicity, New Hampshire. - Emerg. Infect. Dis. 15: 585–587.

Walters, W. et al. 2015. Improved bacterial 16S rRNA gene (V4 and V4-5) and fungal internal transcribed spacer marker gene primers for microbial community surveys. - mSystems 1: e0009-15.

Wang, G. et al. 1997. Genetic and Phenotypic Analysis of Borrelia valaisiana sp. nov. (Borrelia Genomic Groups VS116 and M19). - Int. J. Syst. Bacteriol. 47: 926–932.

Wang, Q. et al. 2007. Naive Bayesian classifier for rapid assignment of rRNA sequences into the new bacterial taxonomy. - Appl. Environ. Microbiol. 73: 5261–5267.
